# Positive effects of landscape diversity on vegetation productivity at the kilometer scale

**DOI:** 10.1101/2025.09.25.678498

**Authors:** Simon Landauer, Florian Altermatt, Roman Flury, Reinhard Furrer, Forest Isbell, Jonas I. Liechti, Eva Spehn, Pascal A. Niklaus

## Abstract

Broad evidence suggests a positive relationship between local species richness and ecosystem functioning ^1,2^. However, Earth’s terrestrial surface is formed by a mosaic of different habitats, which constitutes a higher level of diversity known as landscape diversity. Recent studies indicate that landscape diversity can also promote ecosystem functions, including primary productivity, but the underlying mechanisms and relevant spatial scales remain poorly understood ^3,4^. Here, we combine satellite remote sensing of vegetation productivity with high-resolution land cover data to investigate the effects of landscape diversity on ecosystem functioning across different spatial scales. Using a plot network covering North America, we find that the observed landscape-wide effects cannot be explained by local-scale interactions along ecosystem interfaces. In a subsequent, global-scale analysis, we instead demonstrate that landscape diversity effects are most strongly associated to landscape diversity at a spatial scale of a few kilometers. This association holds across continents and climatic zones, indicating that this pattern is universal. Our investigation therefore suggests that effective land management and biodiversity conservation strategies must integrate both local species richness and higher-level organizational diversity, such as landscape diversity, to ensure robust ecosystem functioning.

## Main

Biodiversity-ecosystem functioning experiments have demonstrated that more species-diverse plant communities on average are more productive ^5–7^. Similar positive correlations between plant diversity and productivity are often also found in observational studies across ecosystems of a certain type (e.g. grasslands, forests), at least when effects of the environment including climate, nutrient availability, and edaphic conditions, which affect productivity through mechanisms independent of plant species diversity, are accounted for ^8,9^. However, in observational studies the investigated plant communities and ecosystems do not exist independent of their surrounding environment ^10^. Typically, the study plots are embedded in a landscape that resembles a mosaic of different ecosystem types, including heterogeneity in topography, land-uses and species pools.

The non-independence of observational plots from their surroundings has important consequences. For instance, local (α)-species diversity in a plot depends on species dispersal across the ecosystem’s boundary ^11–14^, where the dynamics of meta-populations and meta-communities in a larger landscape can modify local α-species diversity through e.g. anthropogenic influence ^15^. Such dynamics are not necessarily a problem for the quantification of the effects of α-species diversity on plot-level productivity, provided that the mechanisms driving these effects take place within the plot boundaries. Likely mechanisms as such are facilitation and competition related to resources such as soil nutrients and light. However, other mechanisms known to shape diversity-productivity relationships ^16–18^ operate on much larger scales. One example is the epidemiological dynamics of airborne fungal pathogens or insect pest species. These can be transmitted across plot boundaries, with adjacent ecosystems potentially serving as reservoirs for the pest, and thus increasing pest loads within the plot. Alternatively, a heterogeneous landscape can promote the local abundance of pollinators, a phenomenon often observed in agricultural fields that benefit from pollinators that find habitat in nearby more natural ecosystems ^19^. Adjacent ecosystems also exchange matter (e.g., carbon, nutrients, water) and energy (e.g. heat), which can improve resource availability and thereby benefit adjacent ecosystems ^16,18^. Consequently, diversity and spatial arrangement of ecosystem types *per se* may affect functions of larger landscapes such as primary production ^20^.

Studies have tested whether diversity of ecosystem type affects landscape wide functioning and found that remotely-sensed vegetation productivity of landscape patches increased with landscape diversity, both in Switzerland ^3^ and in North America ^4^. The effect of landscape diversity, which was measured as land cover (LC) type diversity, was at least in part independent of additional, positive, effects of the α-species richness of individual ecosystems within these landscapes ^3,4^. These studies provided evidence that ecosystem type diversity is positively associated with landscape productivity across large areas of land. However, the particular mechanisms that drive this positive correlation between landscape diversity and productivity remain elusive, and the spatial scale at which these effects occur unclear. At the lower end of the scale, these diversity effects could originate from interactions that occur along the immediate interfaces between ecosystems ^21^. For example, forest productivity typically is higher at forest edges than in forest interiors ^22^. At the higher end of the scale, effects could also emerge from interactions among ecosystem types that occur at scales much larger than the extent of the landscape patch investigated.

Here, we studied the spatial scale of landscape diversity effects on vegetation productivity, quantified using time-series of satellite-sensed vegetation productivity indices. In a first step, we tested whether LC-type diversity effects originate from interactions along or near the immediate interfaces between different LC types. Specifically, we analyzed productivity data from 36,609 study landscapes (plots) that were spread across most of North America. These analyses were either performed using the entire 500 × 500 m landscape area, or excluding a buffer area along LC interfaces. We speculated that effects predominantly driven by LC-type interactions occur at a spatial range of up to a few ten meters and should decrease or vanish when the area along the LC interfaces is not considered in the productivity estimation of the respective LC patch. In a second step, we tested for LC-type diversity effects that occur at much larger spatial scales. This analysis was performed by convolving a global LC map with Gaussian kernels varying in size (a process we here refer to as spatial “diffusion”), and analyzing landscape productivity using this diffused LC-type information. The underlying rationale was that the productivity at any particular location is affected by all surrounding locations whose Gaussian kernel extend to this location. In other words: locations with a different LC can interact, and thereby potentially produce LC diversity effects, in the area in which their Gaussian kernels overlap. Determining the kernel size (diffusion range) that best explains the observed diversity effects then indicates the likely scale of the underlying LC interactions.

## Results AND Discussion

### Landscape diversity

Larger landscapes harbor diversity at many organizational levels ranging from individuals within species, to species, to communities and ecosystems ^20^. While most functional biodiversity research has focused on the importance of species diversity, we here focus on the diversity of larger contiguous units of land. These «land units» ^23^ are the discernible units that compose the typical mosaic as which landscapes are often perceived. In our study, diversity of land units within these mosaics is operationally quantified by the diversity of distinct land cover (LC) types. This diversity metric is an approximation to ecosystem-type diversity, similar to how plant functional types aggregate functionally similar species, and functional-type diversity therefore approximates the overall functional diversity present among species. Because terrestrial landscapes often are anthropogenically influenced, and non-natural LCs such as crop fields and urban areas interact with their surroundings, anthropogenically-shaped LCs were included in the analysis.

### Local-scale diversity effects

To investigate local landscape-diversity effects, we capitalized on a quasi-experimental study design established earlier ^4^. Our design consisted of 25-ha study plots (500 × 500 m “landscapes”) across North America, with plots containing a land cover type richness (LCR) of 1 to 4. To control for large-scale environmental variation, we defined separate experimental regions (spatial blocks) as the intersection between 14 different ecoregions ^24^ and a 3° latitude × 6° longitude grid (Fig. 1; total of 245 blocks). To derive LC types, we used a 30 m resolution land cover map by the Commission for Environmental Cooperation ^25^, where we aggregated the original LC classes to 6 classes: agriculture, forest, grassland, shrubland, wetland, and urban areas. To avoid confounding effects of the experimental LCR gradient with environmental gradients, we used simulated annealing to determine a subsample all possible plots in which LCR was uncorrelated to major environmental variables. This resulted in a total of 36,609 study landscapes, holding 46 different LC-type compositions. To estimate landscape functioning, we retrieved Normalized Difference Vegetation Index (NDVI) data from Landsat (2008–2022) as a proxy for vegetation productivity ^26,27^. We integrated this data over the growing season (NDVI_GS_), accounting for phenological differences among ecoregion. We then normalized the productivity data by region to account for inherent productivity differences among ecoregions (see Methods). We then calculated the net diversity effect (NE) as NDVI_GS_ of mixed LC landscapes minus the average NDVI_GS_ of corresponding single-LC landscapes within the same block. To test for local-scale diversity effects that occur along LC interfaces, we removed a 40 or 80 m wide corridor along the LC edges of the landscape plots, and calculated net diversity effects as described above.

**Fig. 1.**
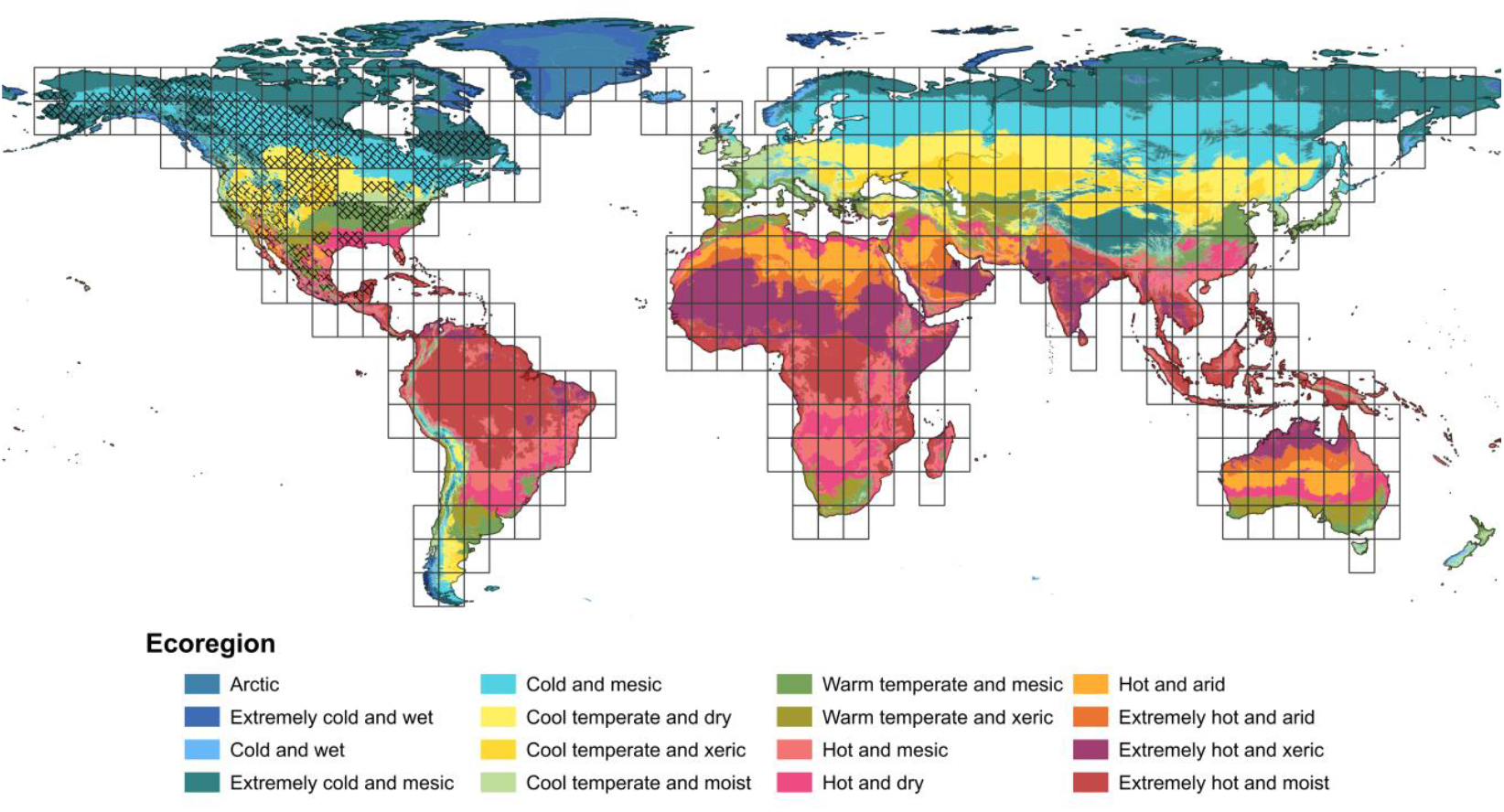
Study area. Landscape diversity effects originating from short range (0 to 80 m) interactions among land cover (LC) types were studied in North America, in 36,609 plots spread across 14 ecoregions (hashed area). Effects occurring at larger ranges (kilometers) were studied at the global scale, in 457 UTM grid zones (rectangles) covering 6 continents and 16 ecoregions.

In all three datasets, LC mixtures were more productive than their corresponding single-LC landscapes (Fig. 2), i.e. net diversity effects (NE) were significantly positive for NDVI_GS_ (P<0.001 for all t-tests and corridor sizes, Table 1). Within LC mixtures, there was a trend for these effects to increase with LCR, at all buffer sizes (P<0.1 for the linear effect of LCR on NE; Fig. 2; Table 1). Overall, diversity-productivity relationships remained virtually unaffected by removal of the area along LC edges, which indicates that the observed LC diversity effects originate at larger spatial scales.

**Table 1.** Net diversity effects on growing-season integrated primary productivity. Net landscape diversity effects were quantified, removing no, 40-m, or 80-m corridor areas along land cover (LC) interfaces. Tests for NE>0 indicate that landscapes containing multiple LC types are more productive that single-LC landscapes. Tests for linear effects o LC type richness (LCR) indicate that there is a marginally significant trend for mixtures with a higher number of LC types to be more productive than less LC-rich mixtures. Both tests did not depend on the extent of the corridor area removed, indicating that the LC diversity effects do not originate from short-distance interactions between LCs along their interfaces. All tests use n=40 different LC composition as unit of replication.

|  | Entire LC area |  | LC area minus 20 m<br>edge buffer<br>(40 m corridor) |  | LC area minus 40 m<br>edge buffer<br>(80 m distance) |  |
| --- | --- | --- | --- | --- | --- | --- |
| NE > 0<br>(across all LC mixtures) | **$t_{39} = 6.20$** | **$P < .001$** | **$t_{39} = 5.98$** | **$P < .001$** | **$t_{39} = 5.77$** | **$P < .001$** |
| Linear effect of LCR<br>(within LC mixtures) | $F_{1, 39} = 3.02$ | $P = 0.09$ | $F_{1, 39} = 3.31$ | $P = 0.077$ | $F_{1, 39} = 3.20$ | $P = 0.081$ |

**Fig. 2.**
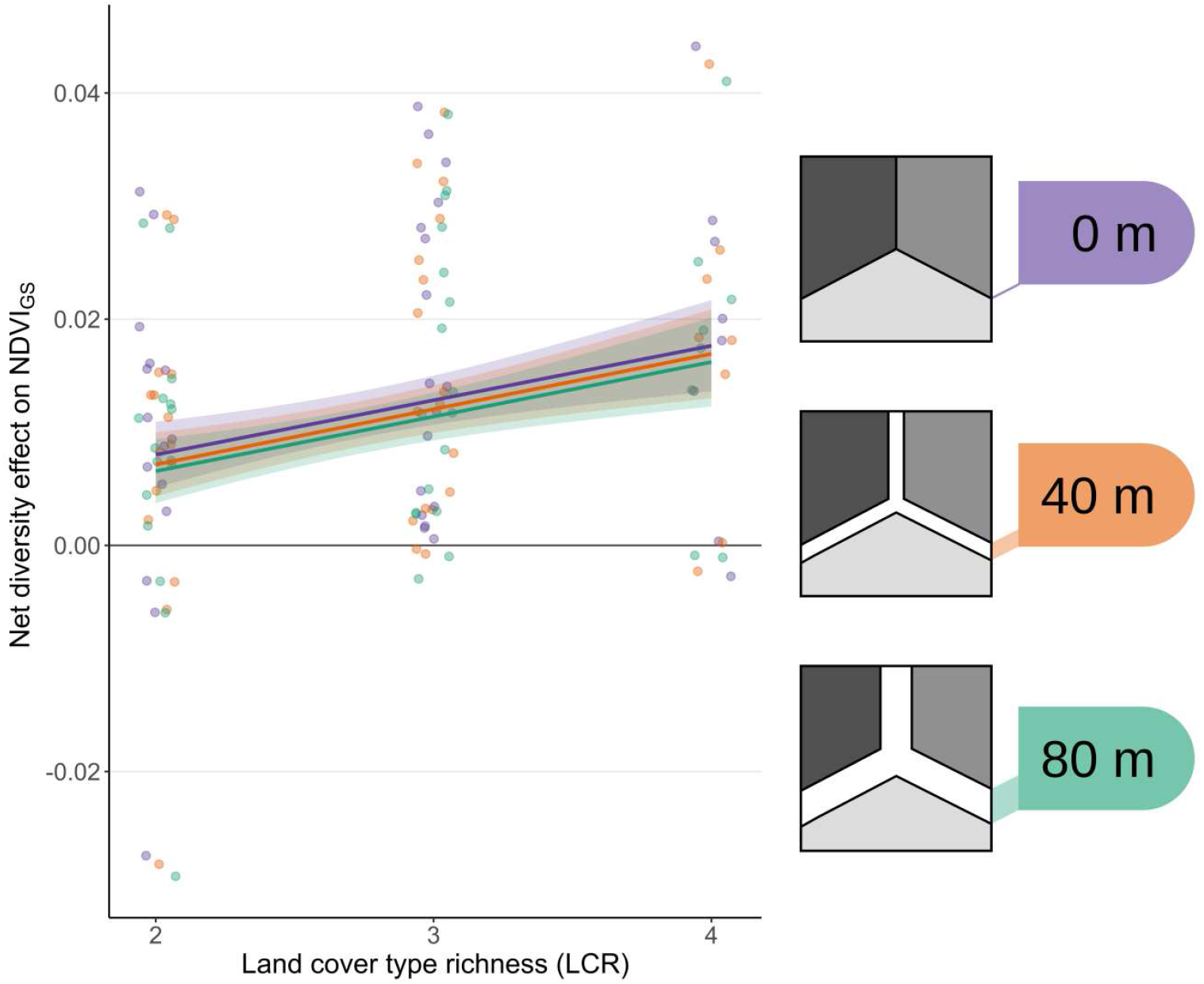
Short-range landscape diversity effects on primary productivity. To test for diversity effect originating from interactions between LC types at a range of up to 80 m, we calculated net effects of LC type richness (LCR) in 500×500 m^2^ plots, removing a buffer between LC types that was 0, 40 or 80 m wide. Primary productivity was quantified as growing-season integrated normalized difference vegetation index (NDVI_GS_). If effects of LCR had been driven by short-range LC-type interactions, the effect of LCR would have decreased with increasing buffer width, which was not the case. Lines and filled areas show model-predicted means and standard errors.

### Large-scale diversity effects

Next, we tested whether the LC-type interactions we were interested may extend beyond the confines of the 500 m landscapes analyzed so far. We reasoned that effects of larger spatial features quantitatively dominate LC interactions, whereas small isolated patches contribute only little as their effect is diluted over comparably large areas. We therefore implemented a spatially-continuous model in which we diffused the LC-effect of each location into its surroundings. Technically, such a diffusion was achieved by convolving a global terrestrial LC map with a two-dimensional Gaussian kernel with unity integral (Fig. 3). We varied the standard deviation of the kernel from ∼42 m to ∼16 km to model LC-diversity effects of different scales (Methods). To isolate large-scale spatial variation, we blocked the study area, this time by the combination of ecoregion with 6° longitude UTM zones × 8° latitudinal bands (Fig. 1). We aggregated LC information from the GLC_FCS30 map (year 2015, 30 m resolution; ^28^) into the eight LC classes cropland, forest, shrubland, grassland, wetland, urban, water, and sparsely or not vegetated. We modeled productivity as additive contributions of the different LC types (the model so far reflecting the null expectation of no LC diversity effects), plus a linear effect of the Shannon diversity index (H), which we calculated based on the diffused LC information.

**Fig. 3.**
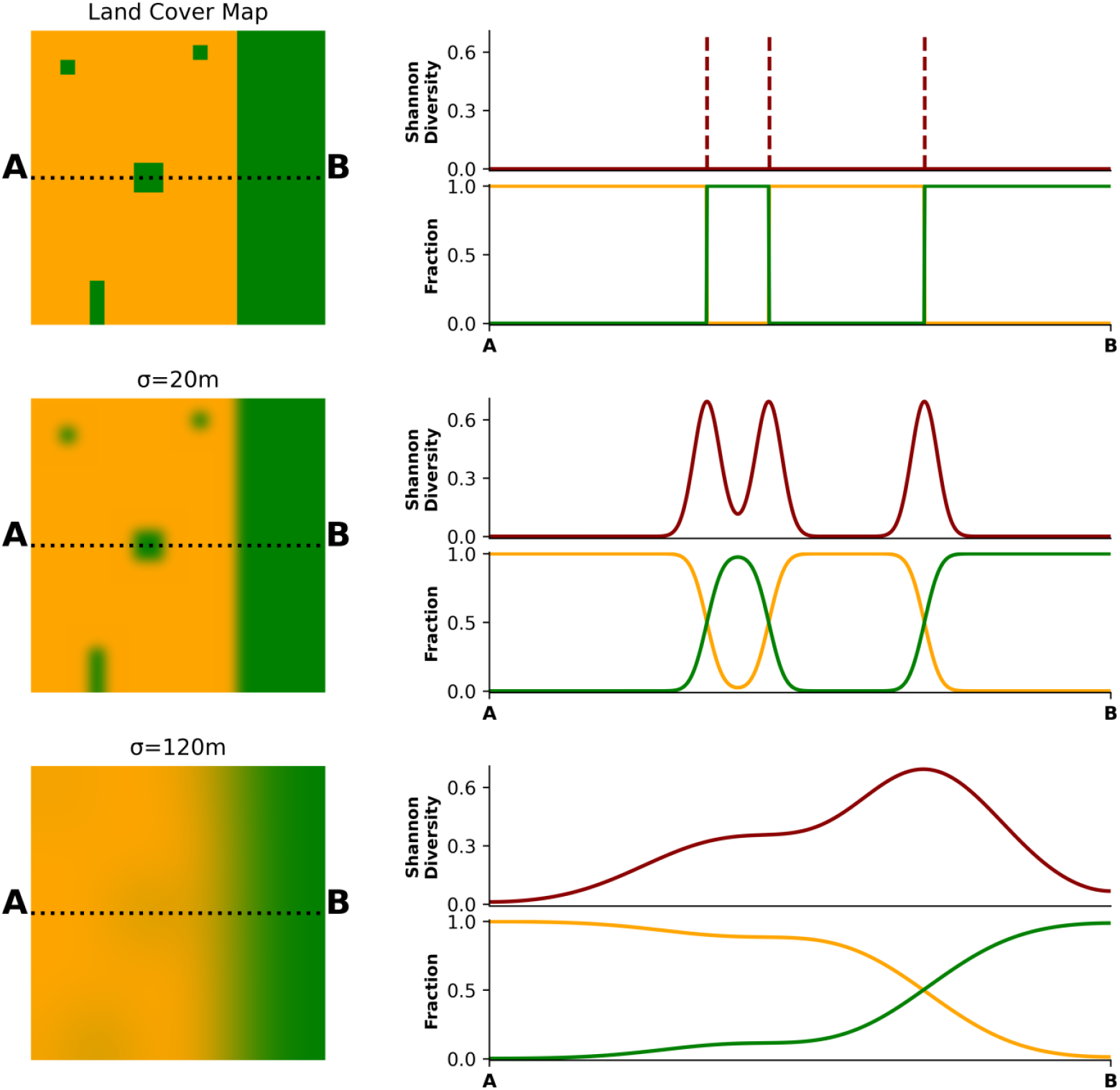
Modeling of long-range landscape diversity effects. To test for effects of land cover (LC) type richness (LCR) occurring at different spatial ranges, LC type information (top left: exemplary 1-km^2^ LC maps with two LC types) was diffused by convolution with a Gaussian kernel with different standard deviations σ (maps on the left, shown here for σ = 20 and 120 m). Then, the Shannon diversity index H_σ_ was calculated based on the diffused LC-type information (right, LC fractions and H along the transect A—B). With infinitesimally small diffusion (σ→0), H equals zero except at the immediate contact area of the two LC types (top right, dashed red line). With increasing σ, the zone in which H substantially exceeds zero becomes wider, and the profile becomes dominated by large LC patches while smaller features disappear.

Comparing models, we found that convolving the LC data with Gaussian kernel sizes with a standard deviation σ of about 1 to 5 kilometers resulted in strongest evidence for positive LC diversity effects (Fig. 4, Table 2), suggesting that the underpinning LC interactions occur over these distances. Given that this analysis covers the entire land area, the LC types are of uneven abundance, with some dominating and some being rarer. LC types also differ in their specific spatial configuration, and as a consequence, different LC types have a different patch size distribution, and have different LCs as their neighbors. The effect of Shannon diversity we determined is thus dominated by the particular LC types and LC type neighborhoods that are abundant in the respective regions, which complicates interpretation.

**Table 2.** Effect of landscape diversity on primary productivity. Landscape diversity was either quantified as Shannon diversity H of land cover (LC) types, or as average pairwise interaction between LC types, based on Simpson’s diversity index D. Primary productivity was determined as normalized difference vegetation index (NDVI) integrated over the growing-season. Diversity indices was based on LC information diffused by convolution with a Gaussian kernel with different standard deviations σ to model different spatial ranges of interactions among LC types. t-tests use the combination of ecoregion and continent as unit of replication (Fig. 1).

| Spatial range $\sigma$<br>(meters) | Shannon diversity | | | Pairwise interaction | | |
| --- | --- | --- | --- | --- | --- | --- |
|  | Landscape<br>diversity<br>effect | t-value<br>(76 df) | P-value | Landscape<br>diversity<br>effect | t-value<br>(89 df) | P-value |
| 42 | 0.0027 | 0.41 | 0.342 | 0.0106 | 2.78 | **0.003** |
| 83 | 0.0021 | 0.29 | 0.387 | 0.0116 | 2.85 | **0.003** |
| 167 | 0.0032 | 0.43 | 0.336 | 0.0129 | 2.99 | **0.002** |
| 417 | 0.0080 | 1.02 | 0.157 | 0.0132 | 2.62 | **0.005** |
| 833 | 0.0148 | 1.85 | **0.034** | 0.0088 | 1.38 | 0.085 |
| 1,667 | 0.0202 | 2.35 | **0.011** | 0.0080 | 0.89 | 0.187 |
| 3,333 | 0.0214 | 2.30 | **0.012** | 0.0080 | 0.69 | 0.247 |
| 5,000 | 0.0197 | 1.98 | **0.026** | 0.0089 | 0.52 | 0.302 |
| 8,333 | 0.0201 | 1.55 | 0.063 |  |  |  |
| 16,667 | 0.0274 | 1.35 | 0.090 |  |  |  |

**Fig. 4.**
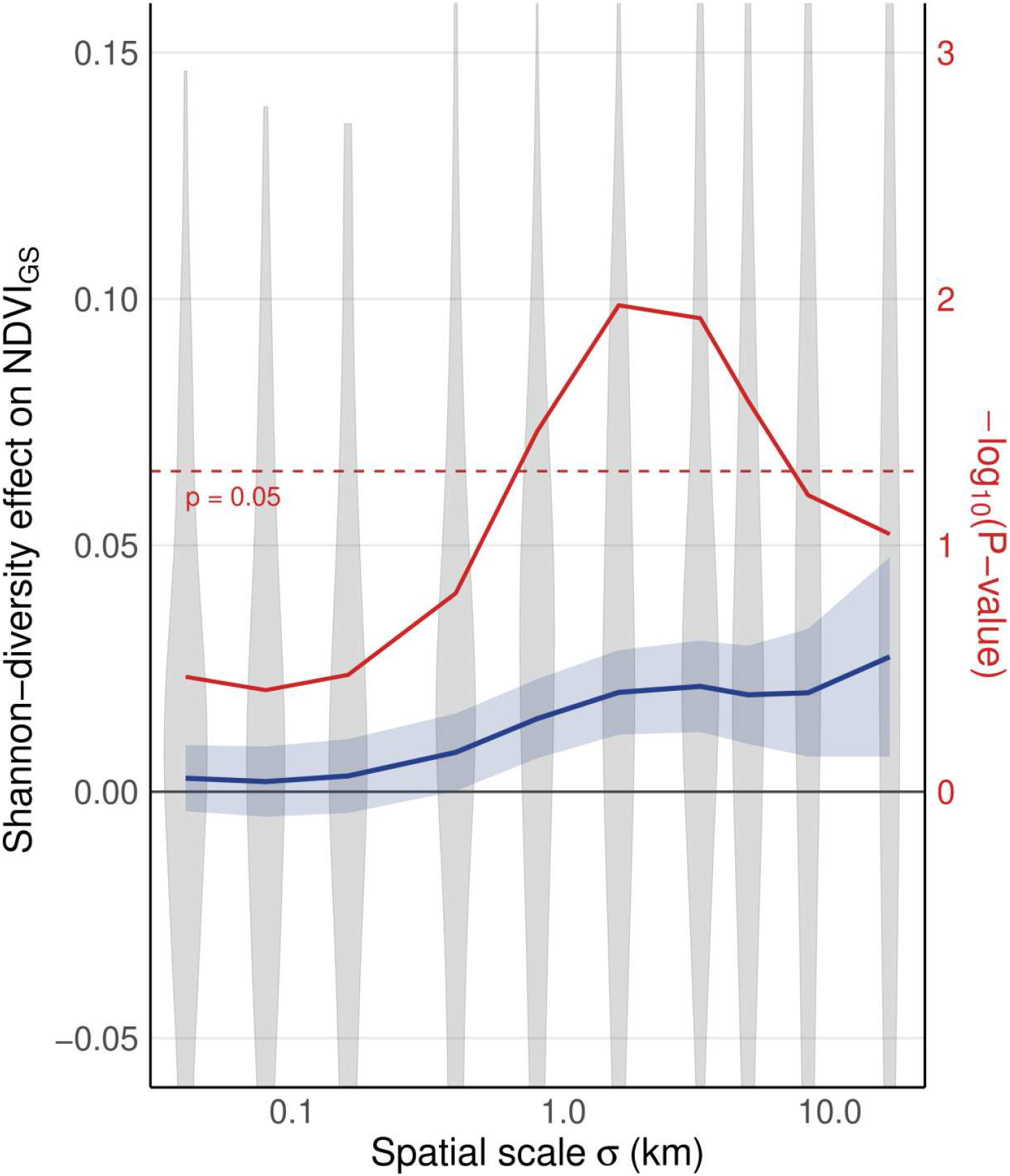
Scale-dependence of effect of Shannon diversity H on primary productivity. Long-range interactions among land cover (LC) types were modeled by diffusing LC-type information with a Gaussian kernel of standard deviation σ (Fig. 3). Primary productivity was determined as growing-season integrated normalized difference vegetation index (NDVI_GS_). Effects were determined as multiple linear regression coefficients of NDVI_GS_ against H (blue line: mean±standard deviation; red line: log_10_ of corresponding P-value) and were statistically significant in the range of a few kilometer. Grey violins indicate the distribution of data at the different levels of σ, using means of ecoregion × continent as unit of replication.

To avoid such potential structural dependency, we next modeled the net diversity effect as sum of all possible pairwise LC-type interactions, instead of using Shannon diversity. Approximating the net diversity effect in this way is justifiable, given that the major productivity gain with LCR occurred between single-LC landscapes and two-LC mixtures, with little additional further increase when LCR was raised beyond two. Given a large enough dataset, pairwise interaction effects are estimable, and, importantly, do not depend on the relative abundance of the LCs in the modeled region. We then averaged the pairwise interaction effects of all possible binary LC combinations, excluding the LC types “water” and “non-vegetated” because these often showed highly negative NDVI values, which cannot reasonably be interpreted as productivity indices. The average NE ultimately obtained is equivalent to determining the diversity effect in a study in which all possible LC combinations are realized at equal replication, as in our quasi-experimental design which balanced the contributions of the different LC types. The average NE was consistently positive for Gaussian kernel sizes σ ranging from a few hundred meters to a few kilometers (Fig. 5, Table 2). Analyzing data by continent demonstrated that this basic pattern is largely universal (Fig. 5).

**Fig. 5.**
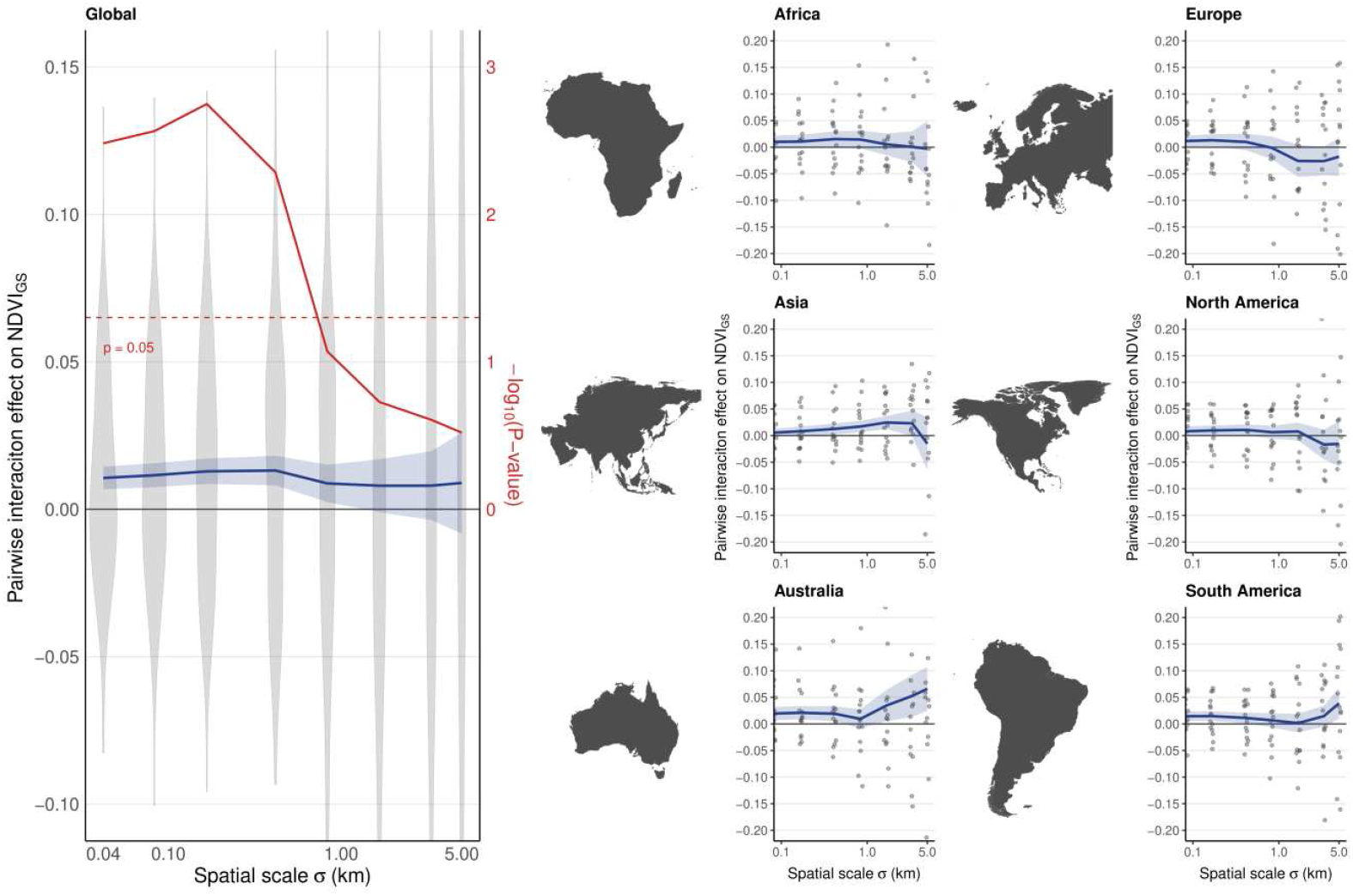
Scale-dependence of pairwise LC interaction effects on primary productivity. Long-range interactions among land cover (LC) types were modeled by diffusing LC-type information with a Gaussian kernel of standard deviation σ (Fig. 3), and determining the average Simpson diversity index D among all LC type pairs present. This effect is independent of the actual abundance of an LC type in a region, i.e. reflects a region in which all pairwise LC-interactions occur at the same frequency. Primary productivity was determined as growing-season integrated normalized difference vegetation index (NDVI_GS_). The effects were determined as multiple linear regression coefficients of NDVI_GS_ against the D (blue line: mean±standard deviation; red line: log_10_ of corresponding P-value), both at the global scale (left) and separately for each continent (right). Grey violins indicate the distribution of data at the different levels of σ, using means of ecoregion × continent as unit of replication.

## Conclusions

Biodiversity-ecosystem functioning research has largely been conducted experimentally on small plots containing a single plant community type. The importance of translating these results to complex real-world landscapes has been recognized ^29,30^, but scaling remains challenging because these landscapes harbor different types of diversity at different scales ^20^, and because it is unclear which level of ecological diversity contributes to landscape functioning and how. Here, we show that composition of LCs that increases LC diversity at the scale of hundreds of meters to a few kilometers increases overall landscape productivity.

Multiple mechanisms may drive these patterns. First, a landscape consisting of a diverse mosaic of different LC types may also be more plant-species-diverse, which may in turn increase productivity. Second, primary productivity may increase because of a parallel increase of beneficial organisms such as pollinators or predators of pests being more abundant in landscapes with more land-use types (thus, species-spill overs across LC types), as is seen in agricultural systems embedded in non-agricultural land-use types ^31,32^. A structurally complex landscape may also confine the spread of pathogens, with benefits possibly extending to non-agricultural ecosystems ^33,34^. Third, abiotic resources, such as nutrients or carbon, may be transferred between ecosystems and subsidize recipient patches/ecosystems ^35,36^. Such transfer includes the airborne movement of volatile nitrogen compounds from an agricultural area to one in which this nutrient is more limiting ^37^. In addition to nutrient transfer, another important resource potentially limiting plant growth is heat. This is evident in growing seasons that are often longer in urban compared to surrounding rural areas ^38,39^. Another example of effects on temperature is found in the woodland-grassland ecotone, which can be warmer due to reduced nighttime cooling in forest ^40^. Whether the mechanisms outlined above indeed underpin landscape diversity effects as the ones we observed is unclear; however the spatial range we have identified may help focus attention on some of these mechanisms.

Many studies have shown that the edges of certain ecosystem types are often more productive than their interior. This effect has been particularly well studied in forests, where enhanced productivity at the edges has been attributed to increased available light, nutrient inputs from surrounding land, and effects resulting from higher species diversity due to increased niche space as consequence of the wider range of environmental conditions ^22,41^. On the contrary, along tropical forests edges higher tree mortality and reduced biomass due to wind and drought were reported ^42^, counterbalancing potential positive edge effects. Interestingly, in the present study, we found no evidence for diversity effects in the immediate vicinity of LC interfaces. One possible explanation is that positive edge effects are manifested only in certain ecosystems (e.g. forests) and are absent or even negative in others, so that across a landscape containing many different ecosystem types, average edge effects are small or absent. Another possibility is that the 30-m LC information used here is too low in spatial resolution to find such edge effects, because productivity at the edges varies at the sub-pixel scale. It is possible that higher productivity at the edge of a forest would then be reflected in a slight increase in the estimated extent of the forest, rather than in increased productivity near the edge. This explanation is consistent with previous research finding that edge length density explained slightly more variance than LC type richness in MODIS satellite-derived NDVI data at 250 m spatial resolution ^3^. In this study, LC information was derived from very high-resolution point grid data obtained from aerial photographs, which minimizes any misattribution of LC type. Overall, it may thus be that in the present study we underestimate effects of LC diversity along edges. Importantly, however, inaccuracy in locating the exact location of LC type-interfaces, if present at all, did not affect our estimates of large-scale LC richness effects, as these remained even when the transition area along the edges was not considered.

Overall, our results highlight the need to better understand the functional consequences of interactions among land uses and land covers at scales larger than interactions along the immediate ecosystem interfaces. To date, effects of forms of biodiversity that emerge at the landscape scale, such as ecosystem type diversity, remain under-explored ^29,30^. Our results indicate that an effective management of ecosystem services might necessitate parallel biodiversity conservation measures at the local and landscape scale, carefully balancing trade-offs between structural diversity and habitat fragmentation.

## Methods

### Test for local-scale landscape-diversity effects

We used a quasi-experimental study design established earlier ^4^ that consisted of 25-ha study plots (“landscapes”) organized in a network that spanned most of North America. As in experimental biodiversity research, we arranged the landscapes in blocks to control for large-scale environmental variation. These blocks were defined by the intersection of 14 ecoregions ^24^ with a 3° latitude × 6° longitude grid (Fig. 1; total of 245 blocks). Within each block, we then established a landscape diversity gradient by selecting landscapes with a land cover type richness (LCR) ranging from 1 to 4. The underlying land cover (LC) types were based on the Commission for Environmental Cooperation’s North American land monitoring system map for 2015 ^25^; 30 m spatial resolution). We aggregated the 19 LC classes of the CEC map into 6 broad, ecologically distinct classes: agriculture, forest, grassland, shrubland, wetland, and urban areas. To keep the LCR gradient independent of selected topographic variables that also affect productivity, we used simulated annealing, a probabilistic subsampling technique. Specifically, we selected a subset of all potential study landscapes (see ^4^ for details) so that elevation and both the north and the east aspect of the slope, which we derived from the TanDEM-X digital elevation model ^43^, were near-orthogonal to LCR. To also keep the average occurrence of each LC type constant along the LCR gradient, we realized all possible LCR combinations within each block, with each combination replicated 20 times. Overall, this design resulted in 36,609 study landscapes and 46 distinct LC type compositions.

As metric of landscape functioning, we used the Normalized Difference Vegetation Index (NDVI), retrieved for the 2008 to 2022 period from Landsat 5, 7, and 8 satellite-sensed multispectral images ^26,27^, using Google Earth Engine ^44^. NDVI data was retrieved separately for each LC found within each landscape, discarding acquisitions with cloud cover over or near the area of interest. Then, growing-season-integrated NDVI values (NDVI_GS_) were obtained by first modeling vegetation phenology as sum of three harmonics ^3^ using the HANTS method ^45^, which is robust to outliers caused by cloud cover and atmospheric disturbance, and then integrating this model from the start (SOS) to the end (EOS) of the growing season. SOS and EOS had previously been determined using the NDVI-ratio method applied to MODIS satellite-derived NDVI data ^4^. Specifically, we used the 25th and 75th percentiles of the estimated start and end of season date samples as SOS and EOS, which results in an interval long enough to accomodate seasonal productivity in areas with above-average growing season length. Next, to be able to compare diversity effects across regions differing in vegetation productivity, we re-scaled NDVI_GS_ values by dividing these by the region’s spatio-temporal NDVI average, which we had determined from an independent large random sample of pixels in each region. Finally, we calculated the net diversity effect (NE) as the NDVI_GS_ value observed in a LC mixture minus the average NDVI_GS_ found in the corresponding single-LC landscapes of the same block.

We repeated the procedure described above for the same set of plots while excluding areas closer than 20m and 40m to LC interfaces, respectively. In other words, this effectively removed a 40 or 80 m wide corridor along the LC interfaces. All data were analyzed using t-tests and linear mixed-effect models fitted with block and LCR as fixed effects and LC composition as random effect ^46^ (VSN International, Hemel Hemsted, UK).

### Test for large-scale landscape-diversity effects

To test for landscape-diversity effects at spatial scales exceeding the 500 m extent of the study landscapes of the quasi-experimental design, a spatially-continuous, global, analysis of vegetation productivity was adopted. We first blocked the global terrestrial surface by ecoregion, and then further into blocks defined by the combination of 6° longitude UTM zones with 8° latitudinal bands (Fig. 1). These blocks correspond to the Military Grid Reference System (MGRS). As LC classification, we used the global LC map GLC_FCS30 (year 2015, 30 m resolution; ^28^) which we aggregated to 8 distinct LC classes: cropland, forest, shrubland, grassland, wetland, urban, water, sparsely or not vegetated. Since mountain shadows were often misclassified as water, we excluded all water pixels above 1000 m a. s. l. and on slopes steeper than 2°. Estimating NDVI_GS_ using the HANTS method requires downloading the time-series of Landsat images, which was computationally not feasible at the global scale. We therefore estimated NDVI_GS_ directly on the Google Earth Engine by dividing the growing season into five equal-length intervals, and estimating the median NDVI value of each interval for the 2014 to 2016 period. Using the median instead of the mean results in insensitivity to outliers in a way comparable to the HANTS method. Finally, NDVI_GS_ was estimated as mean value across the five intervals, multiplied with season length (expressed as fraction of a year). NDVI_GS_ calculated in this way yielded essentially the same values as the HANTS method

To test for scale-dependent interactions among LC types, we convolved the original LC maps with a 2D-Gaussian kernel parameterized by its standard deviation σ, with σ ranging from 42 m to 16 km. Then, we calculated the Shannon diversity index H for each 30 × 30 m LC map pixel based on this diffused LC information, the rationale being that the effect of a particular LC reaches beyond its immediate location, and that nearby LCs therefore interact in the zone where the corresponding Gaussian kernels overlap (Fig. 2). We further calculated indices for pairwise LC interactions as D_A,B_ = 2·x_A_·x_B_ / (x_A_+x_B_), where x_A_ and x_B_ are the fractional LCs of the two LC types, after convolution with the corresponding kernel. D corresponds to Simpson’s diversity index, weighed by the combined abundance of the two LC. Weighing by abundance adjusts for the fact that the observable effect on NDVI_GS_ decreases when the two interacting LCs do not cover the entire plot. We then modeled, separately for each ecoregion × block combination and convolution kernel size σ, NDVI_GS_ pixel values as *NDVI*_*GS*_ ∼ ∑ *x*_*i*_β_*i*_ + *H*β_*H*_. Within this forumla, x_i_ represents the diffused fraction values the respective LC type, and β_i_ the respective coefficients to be estimated; the sum term models additive effects of the different LC types, and H·β_H_ model additional diversity-dependent interaction effects. Similarly, we fitted a model in which diversity effects were modeled as sum of all pairwise LC interactions *NDVI*_*GS*_ ∼ ∑ *x*_*i*_β_*i*_ + ∑ *D*_*i,j*_β_*i,j*_. Because the statistical uncertainty of parameter estimates increased strongly for small sample sizes, we dropped model results for datasets covering less than 90 km^2^ (100,000 pixels). We also excluded individual model terms from the model when the number of pixels with non-zero coefficients (operationally defined as > 0.001) were below 100,000 or below 1% of the total number of pixels, whichever was smaller. The estimated parameter showed a fat-tailed distribution across the up to 540 spatial blocks for which we fitted these models. We therefore removed values that were more than four standard deviations (estimated robustly using the ‘mad’ function in R) away from the median. For the final analysis, we kept only the ecoregion × UTM zone combinations that contained more than two LC types, and for which we could obtain parameter estimates for the full set of Gaussian kernel σ values we investigated (UTM zones used per ecoregion and land cover type pairs, Supplementary Fig. 1, Supplementary Fig. 2). Statistical analyses were finally done on an aggregated datasets with ecoregion × continent combinations as replicates.

## Acknowledgments

This project was funded by a grant of the University of Zurich Research Priority Program Global Change and Biodiversity (URPP GCB) to P.A.N. and R. Fu.. Additionally, this project has received funding from the European Union’s Horizon 2020 research and innovation programme under the Marie Sklodowska-Curie (MSC) grant agreement No 847585.

## Author CONTRIBUTIONS

P.A.N., S.L. and R.Fu.. conceived the idea. P.A.N., E.S. and R.Fu. acquired funding. S.L. implemented the study and analysed the data, primarily supported by P.A.N. and J.I.L. S.L. and P.A.N. wrote the first draft. E.S., F.A., F.I., J.I.L, R.Fu., and R.Fl. contributed to the final manuscript.

## Competing INTERESTS

The authors declare no competing interests.

## Data AVAILABILITY

The datasets used in this study are available on Dryad, under DOI https://doi.org/10.5061/dryad.76hdr7t81.

## Code AVAILABILITY

Computer code for analyses will be deposited on Zenodo after acceptance, and made availbe to reviewers upon request.

## SUPPLEMENTARY Information

**Supplementary Fig. 1.**
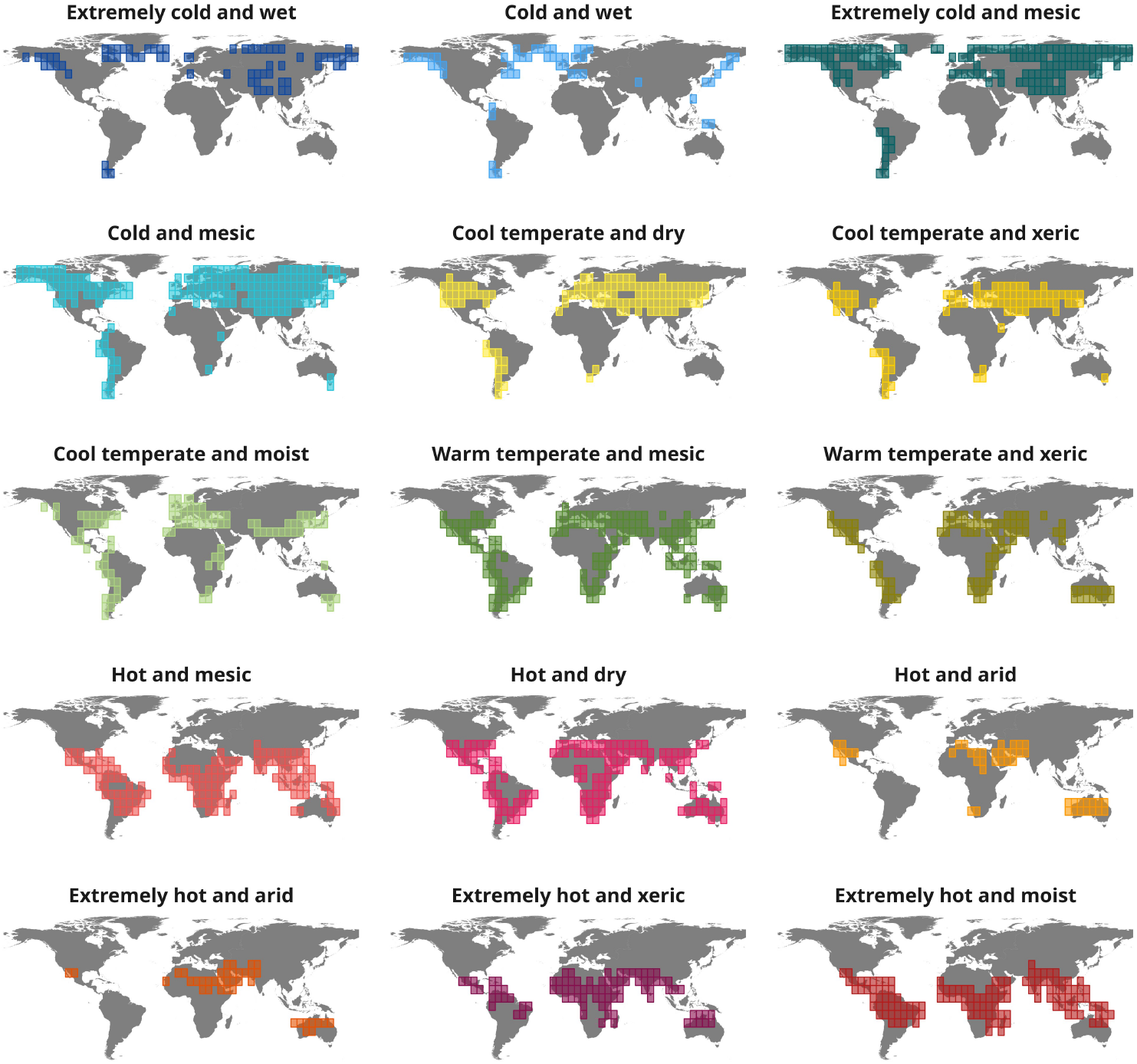
UTM zones underlying data analysis, shown separately for each ecoregion.

**Supplementary Fig. 2.**
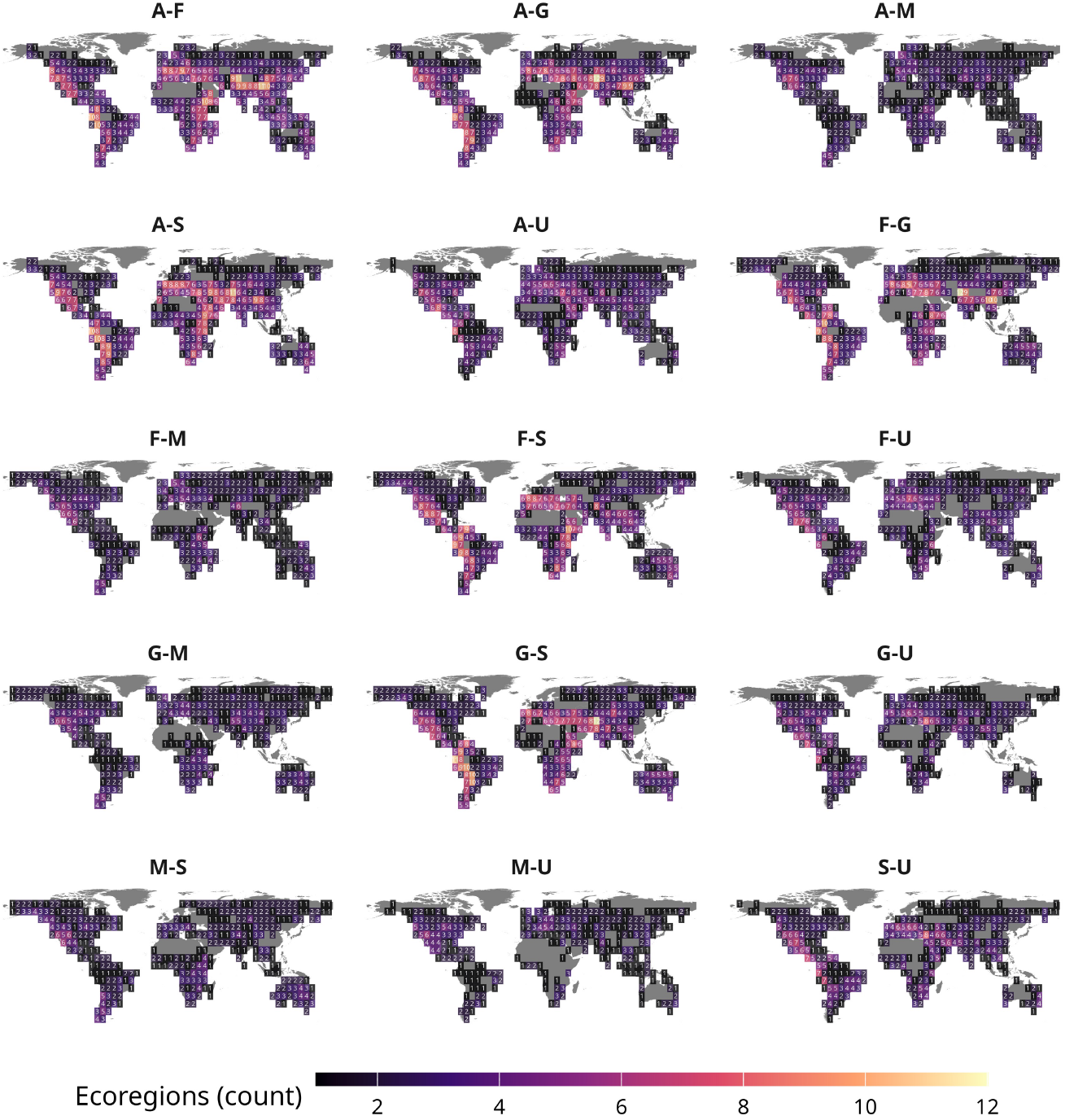
Land-cover type pairs underlying the modeling of diversity effects based on pairwise LC-type interactions. Data are shown at the level of UTM zones, with the color code indicating the number of ecoregions in which the interaction was estimated. A=agriculture, F=forest, G=grassland, M=marshes/wetland, S=shrubland, U=urban.

## Notes

### Competing Interest Statement

The authors have declared no competing interest.

